# Universal target-enrichment baits for anthozoan (Cnidaria) phylogenomics: New approaches to long-standing problems

**DOI:** 10.1101/180067

**Authors:** A.M. Quattrini, B.C. Faircloth, L.F. Dueñas, T.C.L. Bridge, M. Brugler, I.F. Calixto-Botía, D.M. DeLeo, S. Forêt, S. Herrera, S. Lee, D.J. Miller, C. Prada, G. Rádis-Baptista, C. Ramírez-Portilla, J.A. Sánchez, E. Rodríguez, C.S. McFadden

## Abstract

Anthozoans (e.g., corals, anemones) are an ecologically important and diverse group of marine metazoans that occur from shallow to deep waters worldwide. However, our understanding of the evolutionary relationships among the ∼7500 species within this class is hindered by the lack of phylogenetically informative markers that can be reliably sequenced across a diversity of taxa. We designed and tested 16,308 RNA baits to capture 720 Ultraconserved Element loci and 1,071 exon loci. Library preparation and target enrichment was performed on 33 taxa from all orders within the class Anthozoa. Following Illumina sequencing and Trinity assembly, we recovered 1,774 of 1,791 targeted loci. The mean number of loci recovered from each species was 638 ± 222, with more loci recovered from octocorals (783 ± 138 loci) than hexacorals (475 ±187 loci). Phylogenetically informative sites ranged from 26-49% for alignments at differing hierarchical taxonomic levels (e.g., Anthozoa, Octocorallia, Hexacorallia). The percent of variable sites within each of three genera (*Acropora, Alcyonium*, and *Sinularia*) for which multiple species were sequenced ranged from 4.7-30%. Maximum likelihood analyses recovered highly resolved trees with topologies matching those supported by other studies, including the monophyly of the order Scleractinia. Our results demonstrate the utility of this target-enrichment approach to resolve phylogenetic relationships from relatively old to recent divergences. Re-designing the baits with improved affinities to capture loci within each sub-class will provide a valuable toolset to address systematic questions and further our understanding of the timing of diversifications in the class Anthozoa.

## Introduction

Anthozoan cnidarians play critical roles in many marine ecosystems. The class contains ∼7,500 extant species (i.e., soft corals, sea fans, stony corals, black corals, and anemones) that live worldwide in a variety of marine habitats—from tropical shallow waters to the cold deep abyss (Daly *et al*. 2007). Many anthozoans serve as foundation species by creating habitat and supporting a diversity of invertebrates and fishes, including obligate symbiotic associates. In addition, scleractinian (stony) corals can create massive biogenic structures and engineer entire reef-based ecosystems. In fact, coral reefs are some of the most diverse and valuable marine ecosystems across the globe, supporting a high concentration of the oceans’ biodiversity (Carpenter *et al*. 2008). Yet, coral reefs are disappearing at an alarming rate, the health of many coral habitats is declining, and the extinction risk of reef-building corals is increasing due to human impacts, including anthropogenic climate change (Gardner *et al*. 2003; Pandolfi *et al.* 2003; Carpenter *et al*. 2008; Hughes *et al.* 2017). With global ocean change occurring at unprecedented rates (Levitus *et al.* 2000; Gruber *et al*. 2012; Schmidtko *et al*. 2017), it would be helpful to understand the timing of divergence events among particular clades and the history of morphological character (e.g., skeletal type) evolution within the Anthozoa to identify possible connections between those evolutionary events and paleoclimate conditions. However, to advance our understanding of the order and timing of these events, we need to generate a robust, well-resolved phylogeny for the group.

Classification of Anthozoa has traditionally been based on morphological characters such as skeletal morphology, colony organization and soft-tissue anatomy of the polyps (Daly *et al*. 2007), including the arrangement of internal mesenteries (Fautin and Mariscal 1991). Long-standing views have recognized the anthozoan sub-classes Octocorallia and Hexacorallia as reciprocally monophyletic (Daly *et al*. 2007), a view also supported by recent phylogenomic analyses of 10s to 100s of genes (Zapata *et al*. 2015; Pratlong *et al*. 2017). Within each sub-class, however, molecular phylogenetic studies have revealed widespread homoplasy in morphological characters and widespread polyphyly at the ordinal, sub-ordinal, family, and genus levels (e.g., Fukami *et al*. 2008; McFadden *et al*. 2010; Rodríguez *et al*. 2014; Daly *et al*. 2017). Consequently, deep flaws exist in our understanding of the phylogenetic relationships among and within anthozoan orders. Attempts to resolve the deep phylogenetic relationships among anthozoans using molecular data have largely been unsuccessful due to relatively slow evolutionary rates of mitochondrial genomes (Shearer *et al*. 2002; Hellberg 2006; Huang *et al*. 2008; Forsman *et al.* 2009), lack of signal in rDNA (Berntson *et al*. 2001; Daly *et al*. 2003) and difficulty identifying and developing PCR primers for single-copy nuclear genes that can be amplified across the entire class (McFadden *et al*. 2011).

Within most anthozoan orders, there is also a lack of phylogenetic resolution at the species level. This may be due to incomplete lineage sorting in gene trees, insufficient data due to the small number of currently available markers, hybridization, and/or lack of morphological synapomorphies in taxonomy (McFadden *et al*. 2010, 2011, 2017; Prada *et al.* 2014; Rodríguez *et al.* 2014; Grajales and Rodríguez 2016; Daly *et al.* 2017). Currently available markers are insufficient at resolving species boundaries for the majority of anthozoans. For octocorals, an extended mitochondrial barcode (*COI+igr1+mtMutS)* has proven useful for revealing cryptic species and delimiting species boundaries within some clades; however, the divergence criterion proposed (McFadden *et al*. 2011) to elucidate these boundaries is low (>0.5% p-distance) and often no genetic divergence is observed among congeneric species (McFadden *et al*. 2011, Dueñas *et al*. 2014, Pante *et al*. 2015). The low genetic variability in the mitochondrial genome has been attributed to a unique mis-match repair enzyme (*mtMutS)* that potentially repairs mutations (Bilewitch and Degnan 2011) thereby causing reduced mitochondrial sequence variation in octocorals when compared to other metazoans (Shearer *et al*. 2002). Mitochondrial sequence variation is also low in the hexacorals (Hellberg *et al*. 2006; Daly *et al.* 2010), creating difficulties in resolving species boundaries using traditional mitochondrial barcodes (i.e., cox1, COI Barcode of Life, Hebert *et al*. 2003; Shearer and Coffroth 2008). Although several studies have resolved species boundaries using a nuclear ITS marker (e.g., Medina *et al*. 1999; Pinzon and LaJeunesse 2011), using ITS poses problems as it is not a single-locus marker (Vollmer and Palumbi 2004) and there are often high levels of intra-specific variation (Van Oppen *et al.* 2000). Methods that allow for collecting and analyzing hundreds or thousands of loci across shallow and deep levels of divergence are sorely needed.

While there are numerous pathways to find new informative loci (see McCormack *et al.* 2013a), target enrichment of ultraconserved elements (UCEs) (Faircloth *et al.* 2012) has proven robust in inferring species histories of both vertebrates [e.g., fishes (Faircloth *et al*. 2013), birds (McCormack *et al*. 2013b), reptiles (Crawford *et al*. 2012), and mammals (McCormack *et al*. 2012)] and invertebrates [e.g., arachnids (Starrett *et al*. 2016), hymenopterans (Branstetter *et al*. 2017), and coleopterans (Baca *et al*. 2017)] across shallow to deep timescales. UCEs occur in high numbers throughout genomes across the tree of life, including Cnidaria (Ryu *et al*. 2012), making them easy to identify and align among divergent species (Faircloth *et al*. 2012). As the name implies, UCEs are highly conserved regions of the genome, but the flanking regions surrounding UCEs are more variable and phylogenetically informative (Faircloth *et al*. 2012). Some advantages of using target enrichment of UCEs include that 100s to 1000s of loci can be sequenced at a relatively low cost from a wide range of taxa (Faircloth *et al*. 2012); they can be generated from 100 year old, formalin-preserved museum specimens and specimens with degraded DNA (McCormack *et al*. 2016; Ruane and Austin 2017); and they have proven useful at resolving evolutionary questions across both shallow and deep time scales (Smith *et al*. 2013; McCormack *et al*. 2013b; Manthey *et al*. 2016). Similar approaches using target-enrichment of coding regions, or exon-capturing (Bi *et al*. 2012; Ilves and Lopez-Fernandez 2014; Hugall *et al*. 2016), have also proven valuable in phylogenomics.

We used all available genomes and transcriptomes to design a set of target-capture baits for enriching both UCEs and exons for use in anthozoan phylogenetics. Herein, we discuss how loci were targeted and baits were designed. Using an *in silico* analysis, we demonstrate that these loci recover the established sub-class and ordinal relationships among anthozoans. Finally, we test the utility of these baits *in vitro* using 33 species from across both sub-classes of Anthozoa.

## Materials and Methods

### Available Genomes and Transcriptomes

Genomic and transcriptomic data were gathered from various sources for use in bait design and *in silico* testing (Table S1). All data were masked for repetitive regions, retroelements, small RNAs, and transposons using Repeat Masker open-4.0 (Smit *et al*. 2015). The N50 was calculated for each genome using stats.sh in the BBtools package (Bushnell 2015). We then constructed 2bit files for all genomes and transcriptomes (faToTwoBit, BLAT Suite, Kent 2002) and simulated 100 bp paired reads from each genome and transcriptome using the program art_illumina (Huang *et al*. 2012). All programs and parameters used for the entire workflow can be found in Supplemental File 1.

### Identification of UCE Loci and Bait Design

We used the open-source program PHYLUCE (Faircloth *et al*. 2016) and followed the workflow in the online tutorial (http://phyluce.readthedocs.io/en/latest/tutorial-four.html), with a few modifications to identify conserved regions and design baits to target these regions for downstream next-generation sequencing (Faircloth 2017). To find UCEs, we first aligned an average of 34 million simulated reads from each of the four exemplar taxa, *Acropora digitifera, Exaiptasia pallida, Renilla muelleri,* and *Pacifigorgia irene,* to a base genome, *Nematostella vectensis*, using stampy v. 1 (Lunter and Goodson 2011), with a substitution rate set at 0.05. *Nematostella vectensis* (*‘nemve’*) was chosen as the base genome for the primary bait design because it is one of the most well assembled and annotated anthozoan genomes. Approximately 1.3% of the reads mapped to the *nemve* genome; the resulting alignment file was transformed from SAM format into BAM format (samtools, Li *et al*. 2009) and then transformed into a BED formatted file (BEDtools, Quinlan and Hall 2010). These BED files were sorted by scaffold/contig and then by position along that scaffold/contig. We then merged together the alignment positions in each file that were close (< 100 bp) to one another using bedtools. In addition, sequences that included masked regions (>25%) or ambiguous (N or X) bases or were too short (< 80 bp) were removed using phyluce_probe_strip_masked_loci_from_set. These steps resulted in BED files containing regions of conserved sequences shared between *nemve* and each of the exemplar taxa for further analysis. An SQLite table was created using phyluce_probe_get_multi_merge_table, and included 70,312 loci that were shared between pairs of taxa.

We queried the SQLite table and output a list of 1,794 conserved regions shared between *nemve* and the other four exemplar taxa using phyluce_probe_query_multi_merge_table. This list plus phyluce_probe_get_genome_sequences_from_bed was used to extract the conserved regions from the *nemve* genome. These regions were buffered to 160 bp by including an equal amount of 5’ and 3’ flanking sequence from the *nemve* genome. Another filter was performed at this stage to remove sequences < 160 bp, sequences with > 25% masked bases, or sequences with ambiguous bases. A temporary set of sequence capture baits was designed from the loci found in this final FASTA file. Using phyluce_probe_get_tiled_probes, we designed the bait set by tiling 120 bp baits over each locus at 3x density (baits overlapped in the middle by 40 bp). This temporary set of baits was screened to remove baits with >25% masked bases or high (>70%) or low (< 30%) GC content. Any potential duplicates were also removed using phyluce_probe_easy_lastz and phyluce_probe_remove_duplicate_hits_from_probes_using_lastz. Bait sequences were considered duplicates if they were ≥50% identical over ≥50% of their length.

The temporary bait set (2,131 baits, targeting 1,787 loci) was aligned back to *nemve* and the four exemplar taxa using phyluce_probe_run_multiple_lastzs_sqlite, with an identity value of 70% (the minimum sequence identity for which a bait could be an accepted match to the genome) and a minimum coverage of 83%. From these alignments, baits that matched multiple loci were removed. We then extracted 180 bp of the sequences from the alignment files and input the data into FASTA files using phyluce_probe_slice_sequence_from_genomes. A list containing 710 loci shared between at least three of the taxa was created. Based on this list of 710 loci, the anthozoan UCE bait set was re-designed to target these 710 loci using phyluce_probe_get_tiled_probe_from_multiple_inputs, *nemve,* and the four exemplar genomes. Using this script, 120-bp baits were tiled (3X, middle overlap) and screened for high (>70%) or low (< 30%) GC content, masked bases (>25%), and duplicates. This bait set included a total of 5,459 non-duplicated baits targeting 710 anthozoan loci. All above methods were repeated to add more baits to produce additional octocoral-specific baits and capture octocoral-specific loci. We re-ran the analyses using *R. muelleri* as the base genome and *P. irene, Paragorgia stephencairnsi,* and *Antillogorgia bipinnata* as the exemplar taxon to add 1,317 baits targeting an additional 168 UCE loci to the dataset.

### Identification of Exon Loci and Bait Design

To design baits to target exon regions, the above methods were repeated using available transcriptome data. An average of 7 million reads from five exemplar transcriptome-enabled taxa (*A. digitifera,* Cerianthidae*, Edwardsiella lineata*, *Gorgonia ventalina,* and *Paramuricea* sp.) were simulated and an average of 5.8% of these reads per species were aligned to the *nemve* transcriptome. After we converted the alignments to BED files, merged overlapping reads, and filtered data for short loci and repetitive regions, 44,215 conserved sequences were added to an SQLite database. We queried this database and selected 3,700 loci that were shared between *nemve* and the additional five exemplar taxa. Following a second screening for masked regions, high/low GC content, and duplicates, a temporary exon bait set (5,661 baits) targeting 3,633 exon loci was designed. The temporary baits were re-aligned to the transcriptomes of *nemve* and the additional five exemplar anthozoans to ensure we could locate the loci. A set of 906 loci that were shared by *nemve* and the additional five exemplar anthozoans were added to an SQLite database. We re-designed the exon bait set to target these 906 exon loci using phyluce_probe_get_tiled_probe_from_multiple_inputs, *nemve,* and the five exemplar transcriptomes. This bait set included a total of 8,080 non-duplicated baits targeting 906 loci across all anthozoans. To add more octocoral-specific baits and loci, we then repeated the methods with *Paramuricea* sp. as the base transcriptome and *Anthomastus* sp., *Corallium rubrum, Eunicea flexuosa, G. ventalina,* Keratoisidinae sp., and *Nepthyigorgia* sp. as the exemplar taxa to add 4,914 baits targeting an additional 407 loci to the dataset.

### Final Bait Screening

All of the bait sets were screened against one another to remove duplicates (≥50% identical over >50% of their length), allowing us to create a final non-duplicated Anthozoa bait set. We also screened these baits (70% identity, 70% coverage) against the *Symbiodinium minutum* genome by using phyluce_probe_run_multiple_lastzs_sqlite and phyluce_probe_slice_sequence_from_genomes and removed loci that matched the symbiont. Bait names in the final bait FASTA file begin with ‘uce-’ if designed using genomes to target UCEs and ‘trans-’ if designed using transcriptomes to target exons.

### In Silico Test

*In silico* tests were performed to check how well the designed baits aligned to existing genomes and transcriptomes. First, phyluce_probe_run_multiple_lastzs_sqlite was used to align the UCE baits to the nine 2-bit formatted genomes and an outgroup genome (*Hydra magnipapillata)* and the exon baits to the 24 2-bit formatted transcriptomes (Table S1). An identity value of 50% was chosen for alignments. For each bait test, the matching FASTA data were sliced out of each genome or transcriptome, plus 200 bp of 5’ and 3’ flanking regions, using phyluce_probe_slice_sequence_from_genomes. This resulted in an average of 429 ± 178 SD (44 to 599 per species) UCE loci and 497 ± 230 SD (206 to 857) exon loci per anthozoan species (Table 1). To do a final screen for duplicates, loci were matched to baits using phyluce_assembly_match_contigs_to_probes, with a minimum coverage of 67% and minimum identity of 80%. Here, an average of 355 ± 166 SD (25 to 529 per species) non-duplicate UCE loci and 354 ± 210 SD (106 to 670) non-duplicate exon loci were recovered per anthozoan species (Table 1). Each locus was exported into a FASTA file and aligned with MAFFT (Katoh et al. 2002) using phyluce_align_seqcap_align with default parameters.

**Table 1.** Number of loci recovered from *in silico* analyses after initial and final screens for potential paralogs. Also included are the N50 and number of scaffolds for each genome/transcriptome used in analyses.

| Species | #Scaffolds | N50 | # Loci Recovered |  |
| --- | --- | --- | --- | --- |
|  |  |  | Initial Screen | Final Screen |
| Genomes |  |  |  |  |
| *Acropora digitifera* | 4,765 | 191,489 | 462 | 395 |
| *Acropora millepora* | 12,559 | 181,771 | 511 | 414 |
| *Antillogorgia bipinnata* | 426,978 | 3,212 | 230 | 134 |
| *Exaiptasia pallida* | 4,312 | 442,145 | 518 | 417 |
| *Nematostella vectensis* | 10,804 | 472,588 | 496 | 421 |
| *Pacifigorgia irene* | 183,211 | 2,323 | 547 | 491 |
| *Paragorgia stephencairnsi* | 700,190 | 1,793 | 453 | 371 |
| *Renilla muelleri* | 4,114 | 19,024 | 599 | 529 |
| *Stomphia* sp. | 479,824 | 948 | 44 | 25 |
| *Hydra magnipapillata* | 126,667 | 10,113 | 449 | 99 |
| Transcriptomes |  |  |  |  |
| *Acropora digitifera* | 36,780 | 1,575 | 857 | 620 |
| *Acropora hyacinthus* | 67,844 | 422 | 392 | 296 |
| *Anemonia* sp. | 14,279 | 703 | 235 | 106 |
| *Anthomastus* sp. | 9,368 | 610 | 339 | 272 |
| *Anthopleura elegantissima* | 142,934 | 1,489 | 364 | 207 |
| *Cerianthidae* sp. | 12,074 | 646 | 336 | 157 |
| *Corallium rubrum* | 48,074 | 2,470 | 734 | 606 |
| *Edwardsia lineata* | 90,440 | 1,035 | 841 | 623 |
| *Eunicea flexuosa* | 165,709 | 1,095 | 580 | 507 |
| *Exaiptasia pallida* | 60,101 | 2,159 | 553 | 264 |
| *Fungia scutaria* | 155,914 | 1,619 | 290 | 188 |
| *Gorgonia ventalina* | 90,230 | 1,149 | 731 | 670 |
| *Keratoisidinae* | 12,385 | 702 | 541 | 429 |
| *Metridium* sp. | 10,885 | 752 | 222 | 111 |
| *Montastraea cavernosa* | 200,222 | 2,145 | 206 | 128 |
| *Nematostella vectensis* | 27,273 | 1,524 | 836 | 614 |
| *Nephtyigorgia* sp. | 14,677 | 762 | 698 | 619 |
| *Orbicella faveolata* | 32,463 | 1,736 | 408 | 194 |
| *Paramuricea* sp. | 25,189 | 2,645 | 834 | 747 |
| *Pocillopora damicornis* | 70,786 | 976 | 242 | 152 |
| *Porites astreoides* | 30,740 | 661 | 379 | 243 |
| *Protopalythoa variabilis* | 130,118 | 1,187 | 521 | 204 |
| *Scleronephthya* sp. | 8,401 | 683 | 313 | 257 |
| *Platygyra daedalea* | 51,200 | 684 | 483 | 284 |

The resulting alignments were trimmed using GBlocks (Castresana 2000, Talavera and Castresana 2007). We created two final datasets using phyluce_align_get_only_loci_with_min_taxa, in which all locus alignments contained at least 4 of the 10 taxa for the genome data and 9 of the 24 taxa for the transcriptome data. We then concatenated the resulting alignments into separate supermatrices; one containing UCE loci from 10 genome-enabled taxa and the other containing exon loci from the 24 transcriptome-enabled taxa. Maximum likelihood (ML) inference was conducted on each supermatrix using RAxML v8 (Stamatakis 2014). This analysis was carried out using rapid bootstrapping, which allows for a complete analysis (20 ML searches and 200 bootstrap replicates) in one step.

### In Vitro Test

Following the *in silico* test, the list of designed baits was sent to MYcroarray for synthesis. MYcroarray further screened and removed baits that either had repetitive elements or the potential to cross-hybridize (0.007% total baits removed). We then tested the bait set on 33 anthozoan specimens (Table 2), with both sub-classes and all major orders and sub-orders (for Octocorallia) represented.

**Table 2.**
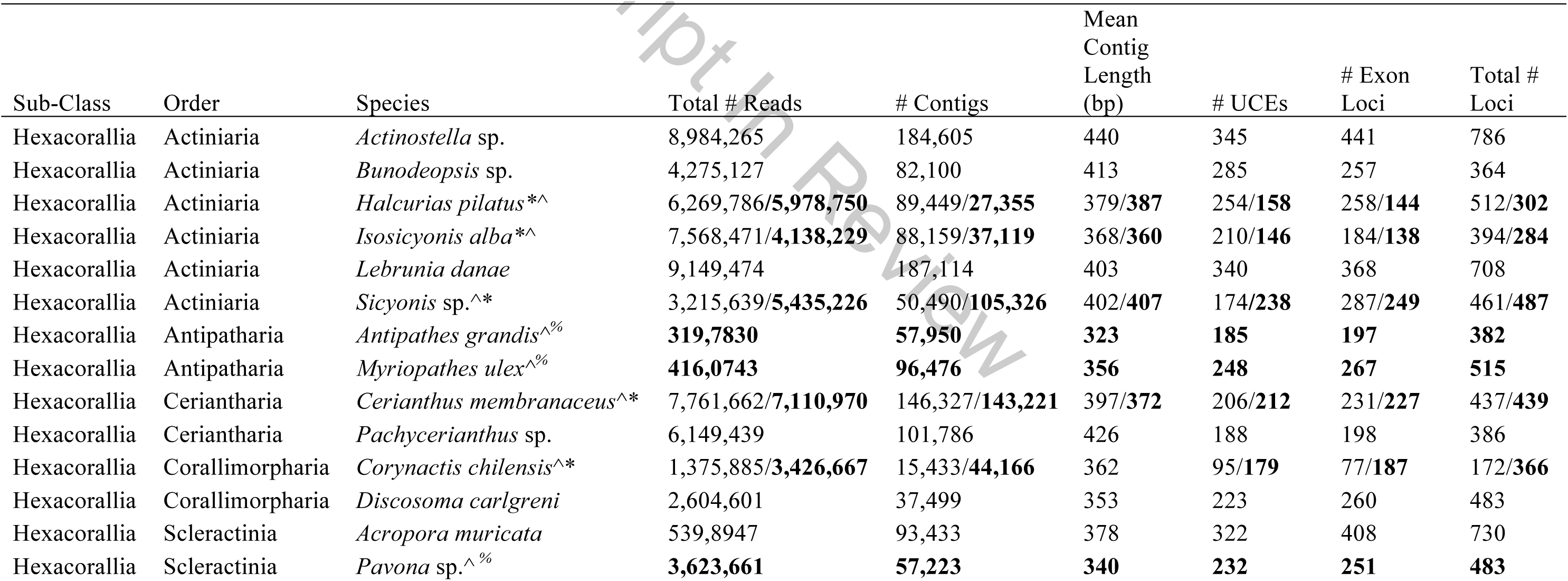
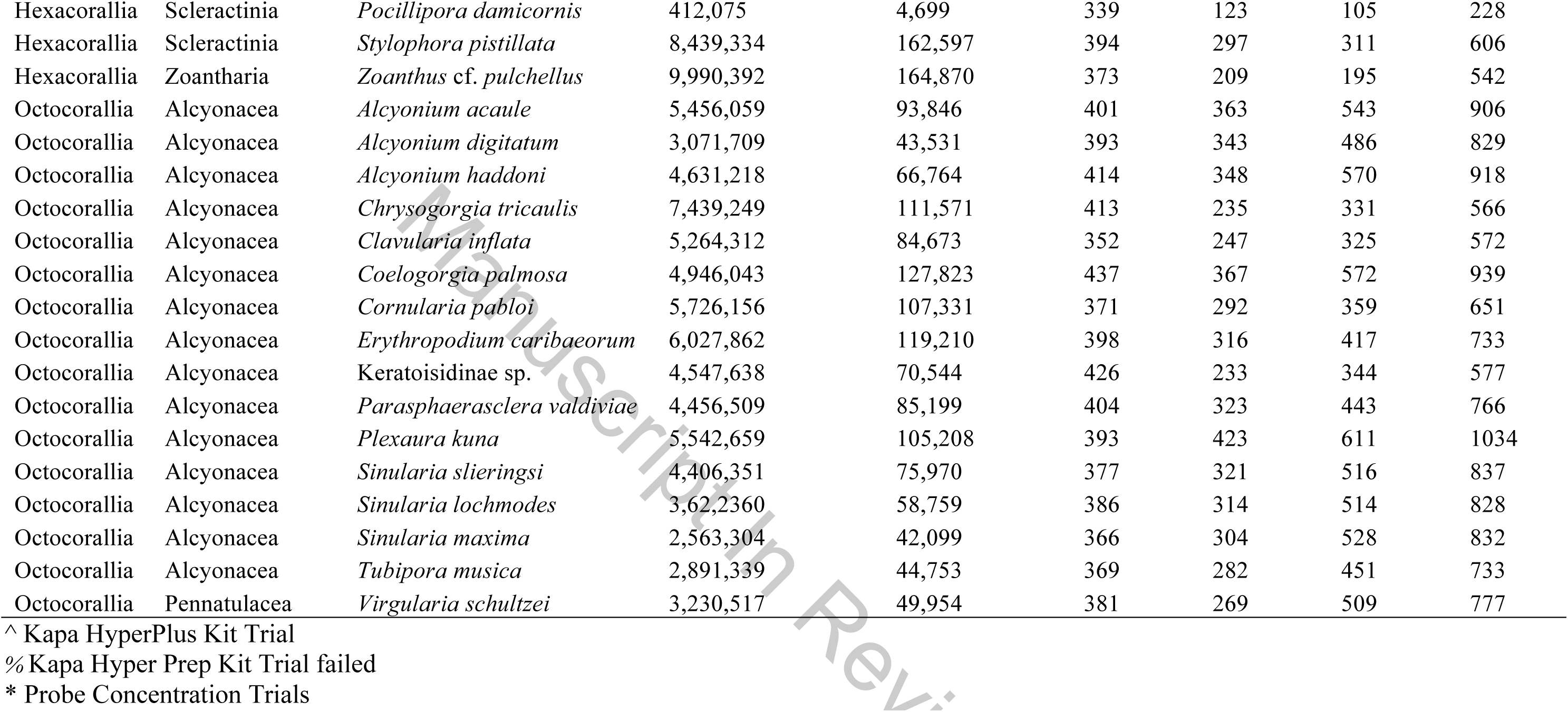
List of species used in the *in vitro* test of designed baits with assembly summary statistics. Results are from the Kapa Hyper Prep and Hyper Plus (in bold) library preparation kits with target enrichments performed using 250 ng of baits.

| Sub-Class | Order | Species | Total # Reads | # Contigs | Mean Contig Length (bp) | # UCEs | # Exon Loci | Total # Loci |
| --- | --- | --- | --- | --- | --- | --- | --- | --- |
| Hexacorallia | Actiniaria | *Actinostella* sp. | 8,984,265 | 184,605 | 440 | 345 | 441 | 786 |
| Hexacorallia | Actiniaria | *Bunodeopsis* sp. | 4,275,127 | 82,100 | 413 | 285 | 257 | 364 |
| Hexacorallia | Actiniaria | *Halcurias pilatus* *^ | 6,269,786/**5,978,750** | 89,449/**27,355** | 379/**387** | 254/**158** | 258/**144** | 512/**302** |
| Hexacorallia | Actiniaria | *Isosicyonis alba* *^ | 7,568,471/**4,138,229** | 88,159/**37,119** | 368/**360** | 210/**146** | 184/**138** | 394/**284** |
| Hexacorallia | Actiniaria | *Lebrunia danae* | 9,149,474 | 187,114 | 403 | 340 | 368 | 708 |
| Hexacorallia | Actiniaria | *Sicyonis* sp.^* | 3,215,639/**5,435,226** | 50,490/**105,326** | 402/**407** | 174/**238** | 287/**249** | 461/**487** |
| Hexacorallia | Antipatharia | *Antipathes grandis*^% | **319,7830** | **57,950** | **323** | **185** | **197** | **382** |
| Hexacorallia | Antipatharia | *Myriopathes ulex*^% | **416,0743** | **96,476** | **356** | **248** | **267** | **515** |
| Hexacorallia | Ceriantharia | *Cerianthus membranaceus*^* | 7,761,662/**7,110,970** | 146,327/**143,221** | 397/**372** | 206/**212** | 231/**227** | 437/**439** |
| Hexacorallia | Ceriantharia | *Pachycerianthus* sp. | 6,149,439 | 101,786 | 426 | 188 | 198 | 386 |
| Hexacorallia | Corallimorpharia | *Corynactis chilensis*^* | 1,375,885/**3,426,667** | 15,433/**44,166** | 362 | 95/**179** | 77/**187** | 172/**366** |
| Hexacorallia | Corallimorpharia | *Discosoma carlgreni* | 2,604,601 | 37,499 | 353 | 223 | 260 | 483 |
| Hexacorallia | Scleractinia | *Acropora muricata* | 539,8947 | 93,433 | 378 | 322 | 408 | 730 |
| Hexacorallia | Scleractinia | *Pavona* sp.^% | **3,623,661** | **57,223** | **340** | **232** | **251** | **483** |
| Hexacorallia | Scleractinia | *Pocillipora damicornis* | 412,075 | 4,699 | 339 | 123 | 105 | 228 |
| Hexacorallia | Scleractinia | *Stylophora pistillata* | 8,439,334 | 162,597 | 394 | 297 | 311 | 606 |
| Hexacorallia | Zoantharia | *Zoanthus cf. pulchellus* | 9,990,392 | 164,870 | 373 | 209 | 195 | 542 |
| Octocorallia | Alcyonacea | *Alcyonium acaule* | 5,456,059 | 93,846 | 401 | 363 | 543 | 906 |
| Octocorallia | Alcyonacea | *Alcyonium digitatum* | 3,071,709 | 43,531 | 393 | 343 | 486 | 829 |
| Octocorallia | Alcyonacea | *Alcyonium haddoni* | 4,631,218 | 66,764 | 414 | 348 | 570 | 918 |
| Octocorallia | Alcyonacea | *Chrysogorgia tricaulis* | 7,439,249 | 111,571 | 413 | 235 | 331 | 566 |
| Octocorallia | Alcyonacea | *Clavularia inflata* | 5,264,312 | 84,673 | 352 | 247 | 325 | 572 |
| Octocorallia | Alcyonacea | *Coelogorgia palmosa* | 4,946,043 | 127,823 | 437 | 367 | 572 | 939 |
| Octocorallia | Alcyonacea | *Cornularia pabloi* | 5,726,156 | 107,331 | 371 | 292 | 359 | 651 |
| Octocorallia | Alcyonacea | *Erythropodium caribaeorum* | 6,027,862 | 119,210 | 398 | 316 | 417 | 733 |
| Octocorallia | Alcyonacea | Keratoisidinae sp. | 4,547,638 | 70,544 | 426 | 233 | 344 | 577 |
| Octocorallia | Alcyonacea | *Parasphaerasclera valdiviae* | 4,456,509 | 85,199 | 404 | 323 | 443 | 766 |
| Octocorallia | Alcyonacea | *Plexaura kuna* | 5,542,659 | 105,208 | 393 | 423 | 611 | 1034 |
| Octocorallia | Alcyonacea | *Sinularia slieringsi* | 4,406,351 | 75,970 | 377 | 321 | 516 | 837 |
| Octocorallia | Alcyonacea | *Sinularia lochmodes* | 3,62,2360 | 58,759 | 386 | 314 | 514 | 828 |
| Octocorallia | Alcyonacea | *Sinularia maxima* | 2,563,304 | 42,099 | 366 | 304 | 528 | 832 |
| Octocorallia | Alcyonacea | *Tubipora musica* | 2,891,339 | 44,753 | 369 | 282 | 451 | 733 |
| Octocorallia | Pennatulacea | *Virgularia schultzei* | 3,230,517 | 49,954 | 381 | 269 | 509 | 777 |

^ Kapa HyperPlus Kit Trial
% Kapa Hyper Prep Kit Trial failed
\* Probe Concentration Trials

DNA was extracted using a Qiagen DNeasy Blood & Tissue kit, Qiagen Gentra Kit, or a CTAB extraction protocol (McFadden *et al*. 2006). DNA quality was assessed using a Nanodrop spectrophotometer, with 260/280 ratios ranging from 1.8-2.1 and 260/230 ratios ranging from 1.4-3.2. The initial concentration of each sample was measured with a Qubit 2.0 fluorometer. For the majority of samples, we then sheared 600 ng DNA (10 ng per μL) to a target size range of 400-800 bp using sonication (Q800R QSonica Inc. Sonicator). For eight samples (Table 2), we sheared 35 μL (115-372 ng, average 217 ng) of EDTA-free DNA using enzymes from the Kapa HyperPlus (Kapa Biosystems) library preparation kit. These samples were mixed on ice with 5 μL of Kapa Frag buffer and 10 μL of the Kapa Frag enzyme and put on a pre-cooled (4**°**C) thermocycler prior to incubation for 10-15 min at 37 **°**C to achieve a target size range of 400-800 bp. After shearing, DNA was run out on a 1% agarose gel (120V, 60 min). Small DNA fragments were removed from each sample (250 ng DNA) using a generic SPRI substitute (Rohland and Reich 2012; Glenn *et al*. 2016) bead cleanup (3X). DNA was re-suspended in 25 μL double-distilled water (ddH20).

Details of library preparation and target enrichment can be found in Supplemental File 2. Briefly, library preparation (Kapa Biosystems) was carried out on the majority of DNA samples (Table 2) using a Kapa Hyper Prep protocol. For the subset of the samples for which DNA was sheared using enzymes (Table 2), we followed the protocol in the Kapa Hyper Plus enzyme-shearing library preparation kit (Kapa Biosystems). Universal Y-yoke oligonucleotide adapters and custom iTru dual-indexed primers were used in library preparations (Glenn et al. 2016). For target enrichment, the MYcroarray MyBaits were diluted in 1/2 (250 ng) of the standard (500 ng) MyBaits reaction, using 2.5 μL of the baits and 2.5 μL of ddH20 for all samples. Different bait strengths were tested on a set of six samples (Table 2): full bait strength (500 ng), 1/2 bait strength (250 ng), 1/4 bait strength (125 ng), and 1/8 strength (63 ng). One pool of enriched libraries was sent to Oklahoma Medical Research Facility for sequencing on 2/3 of a lane of Illumina HiSeq 3000 (150bp PE reads). The remaining 1/3 of the sequencing lane was used for unrelated samples with non-overlapping indexes.

### Post-Sequencing Analyses

De-multiplexed Illumina reads were processed using PHYLUCE following the workflow in the online tutorial (http://phyluce.readthedocs.io/en/latest/tutorial-one.html/), with a few modifications (Suppl. File 1). The reads were first trimmed using the Illumiprocessor wrapper program (Faircloth 2012) with default values and then assembled using Trinity v. 2.0 (Haas *et al*. 2013). We also assembled the data using Abyss 2.0 (Simpson *et al*. 2009) with a kmer value of 31. UCE and exon bait sequences were then separately matched to the assembled contigs (70% identity, 70% coverage) using phyluce_assembly_match_contigs_to_probes to locate the loci. Loci were then extracted using phyluce_assembly_get_match_counts and phyluce_assembly_get_fastas_from_match_counts, exported into separate FASTA files and aligned with default parameters using phyluce_align_seqcap_align, which uses MAFFT. Loci were internally trimmed using GBlocks.

Data matrices of locus alignments were created using phyluce_align_get_only_loci_with_min_taxa, in which each locus had either 25% or 50% species occupancy. Concatenated locus alignments consisted of exon loci only, UCE loci only, and all loci. The number of phylogenetically informative sites was calculated for each alignment across various taxonomic datasets. The script phyluce_align_get_informative_sites was used on the following taxonomic datasets: Anthozoa+genome+outgroup (33 taxa used in *in vitro* test, plus nine genome-enabled taxa and the outgroup *H. magnipapillata),* Anthozoa (33 taxa used in *in vitro* test), Hexacorallia only (17 taxa used in *in vitro* test), and Octocorallia only (16 taxa used in *in vitro* test). The total number of variable sites, total number of phylogenetically informative sites and number of phylogenetically informative sites per locus were calculated. We also calculated the total number of variable sites and the number of variable sites per locus for alignments containing species in each of three genera: *Acropora (A. digitifera, A. millepora, A. muricata), Alcyonium* (*A. acaule, A. digitatum, A. haddoni*), and *Sinularia (S. slieringsi, S. lochmodes, S. maxima).* For the three *Acropora* species, we used loci from one target-capture enrichment sample and from the two *Acropora* genomes that were available.

ML was conducted on each alignment (exon loci only, UCE loci only, and all loci) for the Anthozoa+genome+outgroup taxon set using RAxML v8. This analysis was carried out using rapid bootstrapping, which allows for a complete analysis (20 ML searches and 200 bootstrap replicates) in one step.

## Results

### Identification of Loci and Bait Design

A total of 16,308 baits were designed to capture 1,791 anthozoan loci with 4 to 10 baits targeting each locus. The principal UCE bait set included 5,513 baits designed to target 720 loci. The principal exon bait set included 10,795 baits to target 1,071 loci. Four loci that matched genomic regions in *Symbiodinium minutum* were removed from the dataset. These loci, however, were also detected in azooxanthellate anthozoans, such as *Chrysogorgia tricaulis*.

### In Silico Test

We generated two alignment matrices, one consisting of the exon loci taken from the transcriptome-enabled taxa and the other one consisting of the UCE loci taken from the genome-enabled taxa. The alignment matrix generated with the UCE loci, which included the *H. magnipapillata* outgroup, had a total of 522 loci, with a trimmed mean locus length of 373 bp (95% CI: 8.4) and a total alignment length of 138,778 bp. The alignment matrix generated with the exon loci included 407 loci, with a trimmed mean locus length of 462 bp (95% CI: 5.8) and a total length of 219,339 bp. The ML phylogenies generated from these alignments recovered the previously established sub-class and ordinal relationships within Anthozoa (Fig. 1). The phylogeny generated with the UCE loci had 100% support at all the nodes (Fig. 1a) whereas the phylogeny generated with the exon loci had complete support at the majority (86%) of the nodes (Fig. 1b).

**Figure 1.**
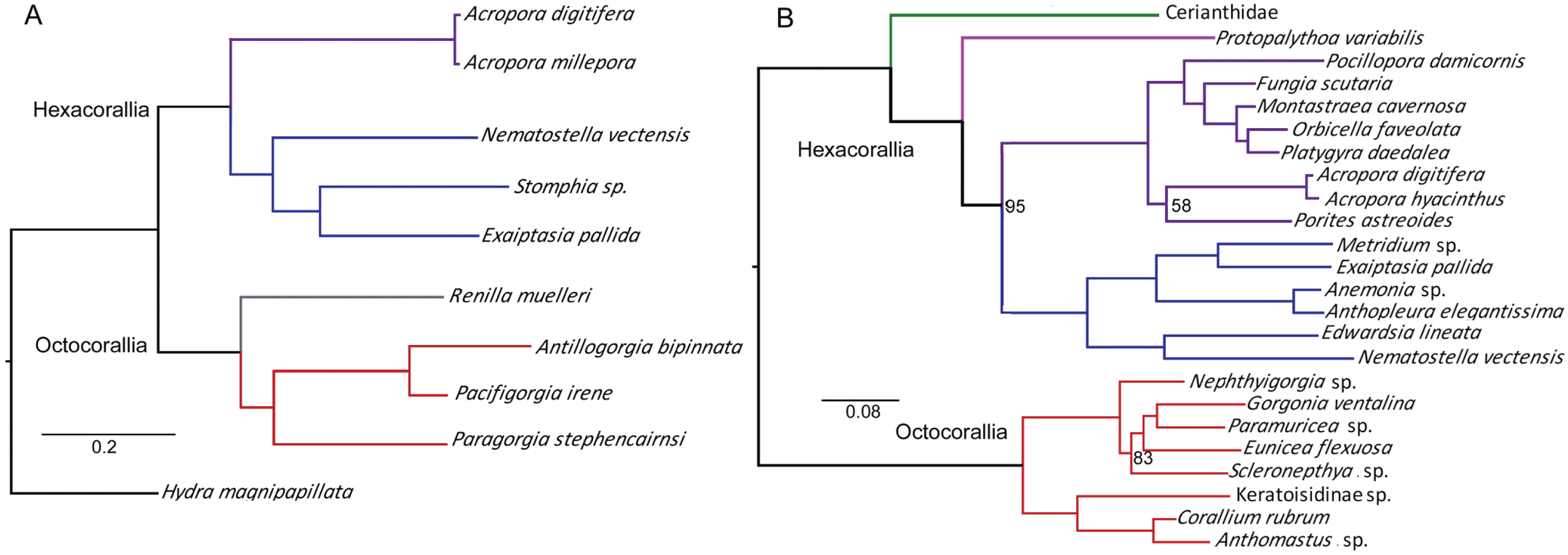
Maximum likelihood phylogenies from *in silico* analyses with bootstrap-support values. A) Phylogeny constructed with a 138,778 bp concatenated genomic dataset (522 loci) and rooted to *Hydra magnipapillata.* B) Phylogeny constructed with 219,339 bp concatenated transcriptome dataset (407 loci). The Hexacorallia clade was rooted to the Octocorallia clade. Nodes have 100% bootstrap support unless indicated. Branches are color coded by order (green=Ceriantharia, pink=Zoantharia, purple=Scleractinia, blue=Actiniaria, red=Alcyonacea, grey=Pennatulacea)

### In Vitro Test

The number of reads obtained from Illumina sequencing ranged from 460,724 to 17,283,798 reads per sample (mean: 5,938,769 ±. 3,407,199 SD reads) across all bait strengths and Kapa kits tested. Quality and adapter trimming lead to the removal of 1.8 to 10.5% reads from each sample, resulting in a mean of 5,486,800 ± 2,092,161 SD trimmed reads per sample (Tables 2, S2, S3). Trimmed reads were assembled into 4,699 to 327,623 contigs per sample (mean: 92,076 ± 65,772 SD contigs) with a mean length of 384 ± 27 bp (range: 224 to 32,406 bp) using Trinity (Tables 2 and S3). Coverage averaged 2.5 to 9.9X per contig. No differences in numbers of contigs or reads were evident between the individuals prepared using the two different Kapa kits (Hyper Prep or Hyper Plus) at 1/2 bait strength or between the different bait strengths used (1/4, 1/8, 1/2, full) (Fig. S1, Tables 2 and S3). Using Abyss, trimmed reads were assembled into 43,428 to 763,227 contigs per sample with a mean length of only 179 ± 24 bp. Because contig sizes were much smaller from Abyss than those assembled via Trinity, remaining analyses were done on the Trinity-assembled data.

A total of 713 UCE loci and 1,061 exon loci (1,774 total loci out of 1,791 targeted UCE loci) were recovered from the assembled contigs. Mean length of UCE contigs was 598 ± 158 bp (range: 224 to 3,995 bp) and mean length of exon contigs was 593 ± 156 bp (range: 224 to 4,500 bp). No differences in numbers of loci recovered were evident between the two different Kapa kits (Hyper Prep or Hyper Plus) at 1/2 bait strength or between the individuals subjected to the four different bait strengths used (Fig. S1, Tables 2 and S3). The number of loci recovered from each species using a Kapa Hyper prep kit with 1/2 bait strength was highly variable, ranging between 172 to 1034 total loci per sample (mean: 638 ± 222 loci) (Tables 2 and S3), although few loci (172) were recovered from the sample with the fewest contigs (15,433). More loci were recovered from octocorals (mean: 783 ± 138 loci, range: 569-1036 loci) compared to hexacorals (mean: 475 ±187 loci, range: 172-786 loci), even after removing the sample with the fewest loci (498 ±172 loci).

Alignment lengths, locus number and length, and the number of phylogenetically informative sites varied depending upon percent (25 or 50%) of taxon occupancy per locus and type of taxonomic dataset (Anthozoa+genome+outgroup, Anthozoa, Hexacorallia, Octocorallia) included in the GBlocks trimmed alignments (Table 3). The average percentage of phylogenetically informative sites across all alignments was 39%. For the comparisons within each of three genera (*Acropora, Alcyonium, Sinularia*), 382 to 426 loci were retained in the 100% alignment matrices (Table 4). Mean % variable sites per locus ranged from 4.7 to 30%, with the most variation found in the *Alcyonium* dataset and the least found within *Acropora*. Percent variation per locus ranged from 0 to 55%, with only one non-polymorphic locus found in the *Acropora* dataset.

**Table 3.** Alignment matrix statistics for different taxonomic datasets. Matrix percentage equals the percent occupancy of species per locus. PI=parsimony informative sites.

| Dataset | % Matrix | # Loci | # Loci (UCE, exon) | Alignment Length | Mean Locus Length (± SD bp) | Locus Length Range (bp) | # PI Sites | %PI Sites |
| --- | --- | --- | --- | --- | --- | --- | --- | --- |
| Anthozoa+genome+outgroup* | 50 | 429 | 228, 201 | 81,403 | 190 ± 89 | 23-549 | 40,041 | 49 |
| | 25 | 1375 | 626, 749 | 257,153 | 187 ± 91 | 23-601 | 119,117 | 46 |
| Anthozoa | 50 | 464 | 229, 235 | 91,455 | 197 ± 93 | 50-667 | 43,501 | 48 |
| | 25 | 1330 | 575, 755 | 254,596 | 191 ± 99 | 19-823 | 109,930 | 43 |
| Hexacorallia | 50 | 438 | 223, 215 | 89,757 | 205 ± 93 | 52-693 | 34,390 | 38 |
| | 25 | 1052 | 529, 523 | 248,476 | 236 ± 107 | 52-1362 | 63,968 | 26 |
| Octocorallia | 50 | 831 | 334, 496 | 208,869 | 251 ± 127 | 51-967 | 70,369 | 34 |
| | 25 | 1366 | 548, 818 | 368,275 | 270 ± 132 | 51-1013 | 96,255 | 26 |

\* includes 33 taxa used in test run, 9 genome-enabled taxa, and the outgroup *Hydra magnipapillata*

**Table 4.**
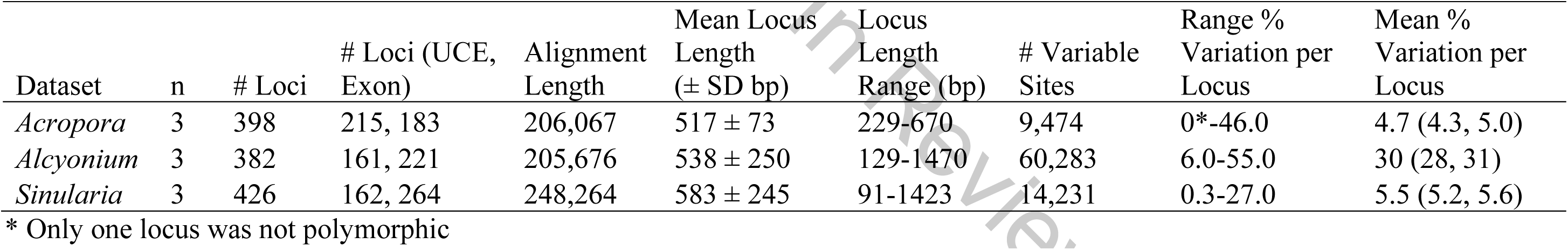
Summary statistics for congeneric species alignments. Mean % variation per locus is also included for UCE loci and exon loci, respectively (in parentheses).

Tree topologies were mostly congruent between the 25% and 50% Anthozoa+genome+outgroup data matrices using all loci (Fig. 2., Fig. S2). By rooting to the outgroup *H. magnipapillata,* monophyly for the currently established anthozoan subclasses and the hexacoral orders was recovered in both taxon occupancy data matrices. As expected, the octocoral order Pennatulacea was found nested within Alcyonacea. Only a few branches shifted between the two data matrices. *Acropora digitifera* was sister to *A. muricata* in the 50% dataset, but sister to *A. millepora* in the 25% dataset. In Octocorallia, both *Cornularia pabloi* and *Erythropodium caribaeorum* shifted positions between datasets. In addition, bootstrap support was higher in the 25% Anthozoa+genome+outgroup tree (Fig. 2) compared to the 50% dataset tree. At most nodes in the 25% dataset tree, bootstrap support was 97-100%; only three nodes had lower bootstrap support (59-82%, Fig. 2).

**Figure 2.**
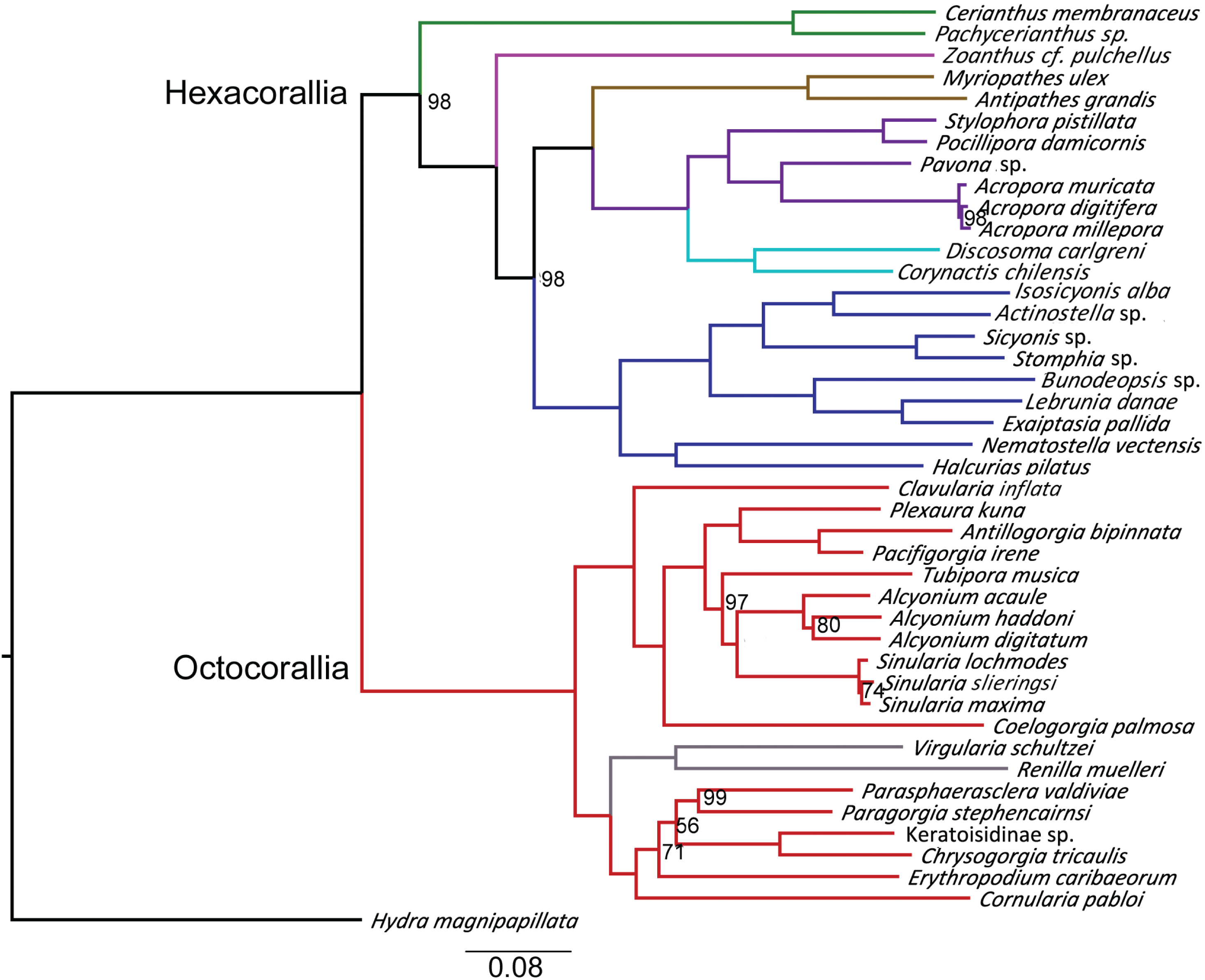
Maximum likelihood phylogeny on the Anthozoa+genome+outgroup 25% matrix (257,728 bp, 1378 loci) with bootstrap-support values. The tree includes 33 taxa from the *in vitro* test, 9 genome-enabled taxa, and the outgroup *Hydra magnipapillata.* Nodes have 100% bootstrap support unless indicated. Branches are color coded by order (green=Ceriantharia, pink=Zoantharia, brown=Antipatharia, purple=Scleractinia, lt. blue=Corallimorpharia, blue=Actiniaria, red=Alcyonacea grey=Pennatulacea)

Lower bootstrap support also occurred in trees created with only the exon loci or the UCE loci (Fig. S3), but tree topologies were mostly congruent with the few exceptions noted above (Fig. S3). We also found that the cerianthids were sister to all other anthozoans in both 25% and 50% exon datasets, but sister to hexacorals in the UCE locus datasets. *Zoanthus* cf. *pulchellus* was sister to the actiniarians in the 25% exon dataset, but sister to a clade containing Actiniaria, Antipatharia, Corallimorpharia, and Scleractinia in all other datasets (Fig S3).

## Discussion

Our results demonstrate the utility of the target-capture enrichment approach for inferring phylogenomic relationships in the class Anthozoa. To date, a few studies based on transcriptomic data have recovered well-supported phylogenomic relationships within Anthozoa, but these studies were based on only a handful (≤15) of taxa (Zapata *et al*. 2015; Lin *et al*. 2016, Pratlong *et al*. 2017) and were limited in scope. In general, phylogenomic studies based on transcriptomic data have provided well-supported and well-resolved phylogenies based on 100s to 1000s of orthologs (Dunn *et al*. 2008; Kocot *et al*. 2011; Zapata *et al*. 2015). However, obtaining these types of sequencing data can be relatively expensive and requires high-quality RNA, two limitations that hinder the transcriptomic-approach for large datasets. In addition, it is often not feasible to obtain RNA from rare taxa or taxa that have not been properly preserved for transcriptomics, such as museum specimens. In our study, we show that the sequence-capture approach for both UCEs and exons can be used to capture genome-scale data in anthozoans. To date, this approach has not been applied to anthozoans or to marine invertebrates more generally (except Hugall *et al*. 2016). We successfully designed a novel bait set based on existing transcriptomes and genomes, and captured 1,774 loci from a diversity of anthozoans spanning >500 million years of divergence (Peterson *et al.* 2004). This target-enrichment approach has the capability to resolve a wide range of divergence levels, from deep-(orders, sub-orders) to shallow-level (species) evolutionary relationships. This novel genomic resource can help to advance studies of systematics, divergence-time estimation, character evolution, and species delimitation in the species-rich class Anthozoa.

### In Vitro Test Results

The newly designed bait set successfully enriched 713 UCE loci and 1,061 exon loci across a diversity of anthozoans. These loci had an average of 39% phylogenetically informative sites, comparable to the arachnid (30% PI sites, Starrett *et al*. 2016) UCE dataset, which targeted ∼1,000 loci. The large range of loci recovered per anthozoan species (172 to 1036 loci) was also similar to the arachnid results (170 to 722 loci). We note that the number of loci recovered from octocorals was much higher than what was recovered from hexacorals. This result is perhaps because we added more octocoral-specific baits to the final bait set. And as we added more octocoral-specific baits, we removed baits that were potential paralogs; the majority of these were designed based on the hexacorals. As was done for the hymenopteran UCE bait set (Branstetter *et al*. 2017), we need to re-design the baitset and include additional octocoral-specific baits and hexacoral-specific baits to increase the success of locus capture. We will also design separate octocoral-and hexacoral-specific bait sets so that additional loci specific to each sub-class can be targeted. Nevertheless, this first bait design and *in vitro* results from 33 taxa demonstrate the promising utility of the target-capture method for resolving anthozoan relationships across deep divergence levels.

The number of variable sites found at loci recovered from within three genera demonstrates that this is also a promising approach to delimit species boundaries. Within all three genera examined, variable sites ranged up to 55% per locus, with a mean variation across all loci of 4.7, 5.5, and 30% in *Sinularia, Acropora*, and *Alcyonium*, respectively. The high variation seen within *Alcyonium* is consistent with unpublished data (C. McFadden, unpubl. data) that suggest the three species are perhaps different genera. For *Sinularia,* average divergence estimates are also higher (∼10X) than what has been demonstrated in other studies using mitochondrial barcoding markers (McFadden et al. 2009). In fact, a 0.5% divergence level at an extended mitochondrial barcode (mtMutS+igrI+COI) was proposed as a conservative criterion for species delimitation (McFadden *et al*. 2011, 2014). Similarly, low divergence estimates at mitochondrial barcoding markers have been found among hexacoral congeners (Shearer and Coffroth 2008; Brugler *et al.* 2013; González-Muñoz *et al.* 2015). Thus, these UCE and exon loci are promising for resolving species boundaries, although we still need to determine the level of intraspecific variation. Recently, restriction-site associated sequencing (RADSeq) has been successfully used to delimit species within anthozoan genera (e.g., Combosch and Vollmer 2015; Pante *et al.* 2015; Herrera *et al*. 2016; McFadden *et al*. 2017; Johnston *et al.* 2017). Although RADSeq is an effective approach for delimiting species and population structure, addressing deeper-level relationships is not feasible due to locus drop out (Althoff *et al*. 2007; McCormack *et al*. 2013a). Our UCE and exon locus datasets may serve as an alternative resource to address species-boundary questions while allowing for data to be combined and examined across deeper levels.

Because this was the first time the target-enrichment UCE approach had been tested on anthozoans, we compared different concentrations of baits and different library preparation kits to determine whether or not particular methods would recover more loci. We found no differences in the number of loci recovered using different concentrations of baits in the hybridization and enrichment protocols. This bait-strength test suggested that the number of hybridizations obtained from one standard reaction could, at least, be doubled. We also found no differences between the two different Kapa kits used. The enzymatic DNA shearing that can be performed with the Kapa Hyper Plus kit may be useful for researchers who do not have access to a sonicator.

Following trimming and aligning of conserved loci, the mean locus length was much shorter (∼190 bp) compared to the mean length of un-trimmed loci (∼600 bp). Therefore, some of the loci included in the ML analyses were relatively short (< 100 bp), particularly in the Anthozoa+genome+outgroup dataset. In alignments between highly divergent taxa (such as between hexacorals and octocorals), numerous poorly aligned positions and divergent positions were filtered with GBlocks. In contrast, the locus size was considerably higher within genera (∼525 bp) because of fewer poorly aligned and divergent positions. Perhaps re-performing the GBlocks internal trimming with less stringent parameters would increase the size of loci in alignments of divergent taxa. Alternatively, by not using GBlocks in the pipeline, we could increase the size of loci and perhaps the accuracy of tree inference (Tan *et al*. 2015). Stringent alignment filtering, as done with GBlocks, can not only increase the proportion of unresolved branches, but can also lead to well-supported branches that are in fact incorrect (Tan *et al*. 2015). Different methods of aligning and filtering data will be explored in future work.

The phylogenies produced from the *in vitro* data were highly supported despite low overall taxon occupancy (>25 or 50% matrices) and inclusion of short loci. Bootstrap support at most nodes was >97% in both trees, although there were a few nodes that had low support and a few branches that shifted between datasets, particularly in the Octocorallia. In addition to stringent filtering as discussed above, sources of incongruence and low bootstrap support could include compositional bias, saturation, violations of model assumptions (Jeffroy *et al*. 2006) and/or missing data. Missing data, however, are generally not problematic if there are a reasonable number of informative characters (see Streicher et al. 2015). Rather, incongruence and low support at a few nodes is perhaps due to incomplete taxon sampling (Wiens 2005; Wiens and Tiu 2012). Although a diversity of taxa from across the clades were selected for *in vitro* analyses, several lineages were not represented, particularly in the Octocorallia. Outgroup choice and taxon evenness can also impact topology and clade support in UCE phylogenomics (Branstetter *et al*. 2017). Future efforts will need to incorporate more thorough taxon sampling.

In general, the inferred phylogenetic relationships corresponded to those found in previous studies (Zapata *et al*. 2015; Rodríguez *et al*. 2014), although there were a few exceptions. One exception was the position of *Cornularia pabloi.* This stoloniferan octocoral was nested within the clade containing sea pens (Pennatulacea) and calcaxonians (*C. tricaulis,* Keratoisidinae sp.), but this species has been found to be sister to the rest of the octocorals based on mitochondrial data (McFadden and vanOfwegen 2012). The superfamily Actinostoloidea (*Sicyonis* sp., *Stomphia* sp.) was recovered as sister to superfamily Actinioidea (*Actinostella* sp., *Isosicyonis alba*) differing from a combined mitochondrial and nuclear rDNA dataset (Rodriguez et al. 2014), which instead recovered Actinostoloidea as sister to both Actinioidea and Metridioidea (*Lebrunia danae, E. pallida*, *Bunodeopsis* sp.). Furthermore, trees in our study were rooted to *H. magnipapillata,* based on the results of Zapata *et al*. (2015); however, the unrooted trees indicated that *H. magnipapillata* was sister to the Octocorallia, a relationship that has been noted in mitochondrial data (Park *et al*. 2012, Kayal *et al*. 2013), but not supported by phylogenomic analyses (Zapata *et al*. 2015). Zapata *et al*. (2015) also found that the position of the order Ceriantharia was phylogenetically unstable. Similarly, our results indicated that the placement of Ceriantharia changed between the different datasets. The topologies resulting from exon data placed the ceriantharians as sister to the anthozoans, a relationship also supported by mitochondrial data (Stampar *et al*. 2014). Trees from UCE loci had ceriantharians as sister to hexacorals, a relationship also supported by combined mitochondrial and nuclear rDNA data (Rodríguez *et al*. 2014). Future work must include different outgroup choices (i.e., sponges), while closely examining the distribution and strength of phylogenetic signal. This will help clarify the source of incongruence and resolve which loci strongly influence the resolution of a given ‘contentious’ branch (Shen *et al*. 2017).

Whether or not scleractinians are monophyletic has been a controversial topic as a result of different phylogenetic analyses. In 2006, Medina *et al*. reported that scleractinians were polyphyletic with corallimorpharians. The “naked coral hypothesis” was thus proposed, suggesting that corallimorpharians arose from a scleractinian ancestor that had undergone skeletal loss during paleoclimate conditions when the oceans experienced increased CO_2_ concentrations (Medina *et al*. 2006). Since that study, other studies based on transcriptomic data (Lin *et al.* 2016), rDNA (Fukami *et al.* 2008), and mitochondrial data (Fukami *et al.* 2008; Park *et al.* 2012, Kayal *et al.* 2013; Kitahara *et al.* 2014) recovered a monophyletic Scleractinia with corallimorpharians as the sister clade. Our results also recovered a monophyletic Scleractinia; thus supporting the conclusions of others that corallimorpharians are not naked corals. However, increased sampling of robust, complex, and basal scleractinians is necessary to conclusively address this issue.

### Future Research Directions

The *in silico* and *in vitro* tests of the novel bait set demonstrate that the target-enrichment approach of UCEs and exons is a promising new genomic resource for inferring phylogenetic relationships among anthozoans. Using this bait set, target-capture enrichment of the UCE and exon loci from at least 192 additional anthozoans is currently underway to further our understanding of character evolution and systematics of the clade. Adding more taxa will likely increase the accuracy of the phylogenetic inference. We also plan to sequence additional outgroup taxa, including medusozoan cnidarians and sponges to help address whether or not octocorals are sister to hexacorals or medusozoans and resolve the position of ceriantharians. Finally, we plan to re-design the bait sets to create hexacoral-and octocoral-specific bait sets.

We will include additional baits to increase the capture efficiency of loci that were targeted in this study, while adding more loci that are specific to each sub-class. This target-enrichment approach provides a promising genomic resource to resolve phylogenetic relationships at deep to shallow levels of divergence, considerably advancing the current state of knowledge of anthozoan evolution.

## Acknowledgements

Funding was provided by NSF-DEB #1457817 to CSM and #1457581 to ER. The Pauley program at the Hawaii Institute of Marine Biology also provided funds to AMQ for preliminary analyses. C. Oliveros and J. Salter helped during lab work and J. Bast helped during preliminary data analyses. F. Zapata and C. Dunn provided transcriptomic datasets. S. Lengyel, E. Cordes, and R. Kulathinal aided DD with the *Paramuricea* transcriptome assembly, which was funded by a grant from the Gulf of Mexico Research to support the “Ecosystem Impacts of Oil and Gas in the Gulf” (ECOGIG) research consortium. E. Bush provided computing support. Special thanks to Z. Forsman, R. Toonen, and I. Knapp for organizing the 2013 HIMB Pauley program.

## Author Contributions

AMQ, CSM, ER, and BCF conceived and designed this study. AMQ designed the baits, conducted library preparation, target enrichment, and data analyses, and wrote the initial draft of the manuscript. BCF developed protocols and guided AMQ in laboratory and bioinformatic analyses. LFD helped with preliminary analyses. MB, ER, and CSM extracted DNA. ICB, DMD, SF, SH, SL, DJM, CP, GRB, CRP, and JAS provided genomic or transcriptomic data for analysis. TB provided samples. All authors edited and approved the final version of this manuscript.

## Data Accessibility

Tree and alignment files: Data Dryad Entry XXXX

Raw Data: SRA Genbank

Anthozoan bait set: Data Dryad Entry XXXX

Scripts: Supplemental file 1

